# Nonmedical Masks in Public for Respiratory Pandemics: Droplet Retention by Two-Layer Textile Barrier Fully Protects Germ-free Mice from Bacteria in Droplets

**DOI:** 10.1101/2020.04.06.028688

**Authors:** Alex Rodriguez-Palacios, Mathew Conger, Fabio Cominelli

## Abstract

Due to the shortage of masks during the pandemic, we recently demonstrated that household textiles are effective environmental droplet barriers (EDBs) with identical droplet retention potential as medical masks. To further promote the implementation of a *universal community droplet reduction solution* based on a synchronized encouragement/enforcement of mask utilization by the public based on widely available textiles (mask fabrication without the need for sewing machines), here we conducted a study using germ-free mice to determine to what extent textiles were effective *in vivo*. Using a bacterial-suspension spray simulation model of droplet ejection (mimicking a sneeze), we quantified the extent by which 100% cotton textile prevented the contamination of germ-free animals on the other side of the textile-barrier (simulating a properly worn mask). Of relevance, all mice protected with textiles remained germ-free after two sprays (inoculation dose: >600 bacterial droplet units per 56.75cm^2^) compared to the contamination of mice not protected by a textile (0/12 vs 6/6, Fisher’s exact, p<0.0001). In a second phase of the experiment with 12 germ-free mice exposed again to 10-fold more droplets remained germ-free, while 100% of mice at 180cm became colonized with a single spray (0/8 vs 4/4, Fisher exact, p=0.002). Collectively, barriers protected all mice (even with low-density textiles, heavy vs. light fabric, T-test, p=0.0028) when using textile-EDB to cover the cages (0/20 vs 10/10, Fisher exact, p<0.0001). This study demonstrated, in vivo, that widely available household textiles are 100% effective at preventing contamination of the environment and the exposed animals by microbe-carrying droplets.

## INTRODUCTION

The economic impact of the COVID-19 respiratory syndrome, declared a pandemic on March 11, 2020, with a doubling time between 2.4 and 5.1 days^1^, will disproportionately affect poor communities^2^. Especially, because lower income individuals have limited resources/access to health-care services, and importantly, because many of these individuals believe that masks are ‘bad’ as they ‘increase the risk of COVID19’, as a consequence of the earlier misleading expert statements and guidelines released to protect the global shortage of medical supplies for hospitals^3–5^. High-exposure risks could also be compounded by limited access to education and income during the crisis^6^ especially among low income ‘lockdown’ communities.

Since COVID-19 transmits primarily via droplet dispersion from symptomatic/asymptomatic individuals as they talk/cough/sneeze^7^, the use of mandatory homemade masks to prevent the contamination of the environment with potentially infective droplets, initially discouraged, has been discussed for voluntary implementation^3–5^. Of interest, the use of masks to prevent droplet dispersion has not been considered as a mandatory strategy, as it has been other measures *(e.g.,* orders to forbid non-essential surgeries^8^) to control COVID transmission by global health directives^9^. At most, some governments started to allow, contradicting initial recommendations, the voluntary use of homemade masks in the community. However, the benefits/implementation of using masks are still debated, with arguments stating that cloths ‘masks increase the risk’. However, such statement is not quantitatively possible if compared to ‘not-wearing masks’.

Because the voluntary use of masks within the community is expected to cause social polarization (believers vs. non-believers; including presidential leaders^10^), if not made mandatory, there is need of further convincing evidence of the ‘mask-wearing’ benefits to incentive their use in public. To prevent the contamination of the environment with COVID-19 droplets, as a continuation of previous studies in masks^11^, and to promote effective education/communication initiatives, herein, we conducted studies using animals born and maintained for life with no germs (germ-free) to determine how effective household textiles are as barriers to protect against microbes inside the droplets.

## METHODS

All respiratory viruses need liquid suspensions/droplets to remain infective for long periods of time (vs. dry), and to contaminate susceptible individuals^12^. Therein, using a bacterial-suspension spray simulation model of droplet ejection (mimicking a sneeze)^11^, and a Parallel Lanes Plating method^13^, herein we quantified the extent by which widely available clothing fabrics could protect germ-free animals on the other side of the textile-barrier (simulating a properly worn mask) from contamination by the microbes contained in micro-droplets. In short, the reported experiment was conducted with eighteen 9-week-old germ-free (Swiss Webster) mice (males:females, 1:1), which were individually allocated to 18 germ-free cages, for repeated exposure to a cloud of micro-droplets.

To test the textile barrier as an effective surrogate alternative simulating a medical mask, we used two layers of a widely common household textile (100% combed-cotton, T-shirts) as cage lid/cover, instead of using the standard germ-free grade mouse cage lids^14^. The choice of textile material was based on their recently proven effect in retaining droplets^11^. Our earlier studies have also shown that two layers of passive filtration are fully protective against viruses/microbes in the room air, when germ-free mice are raised under two-layer of such nested filtration^14^. To determine the extent by which the textile droplet barrier could protect GF mice from droplets, and conduct a statistical powerful study, we exposed to the droplets all animals at a ratio of 2 exposed with EDB:1 without EDB. Animals were observed for three days when fecal cultures were conducted to determine whether animals had been colonized by the bacteria present in the droplet solution used to spray the cages. To further test repeated higher droplet exposure doses, in a second phase of experiment, all animals with EDB that remained germ-free, were exposed again with the textile EDB cover, to 20-sprays (instead of 2; 10-times more droplets) at 60cm, and compared that to animals that were uncovered, and received a single spray-droplet dose at 180cm (minimum social distance recommended; see method details in **Supplementary Materials**).

## RESULTS

Microbiological analysis of the germ-free status of the mice, before and after two rounds of spray-droplet exposure in the first phase of the experiment, showed that all animals after being sprayed with a cloud of droplets at 60cm (inoculation dose: 600-1000 bacterial droplet units per 56.75cm^2^), with no textile protection (simulating not wearing a mask) showed signs of microbial contamination within 18h (fecal culture), by either exposure to the droplets in the environment, or by inhalation, ingestion, or exposure to the droplets on mucous membranes. In contrast, the germ-free status of the mice that were covered with the autoclaved textile EDB, remained germ-free three days after exposure indicating that the textile barrier was extremely effective at retaining bacteria carrying droplets, reducing thus the absolute contamination risk (0/12 vs 6/6, Fisher’s exact, p<0.0001).

The second phase of the experiment (repeated exposure with 10-times more droplets), with 12 germ-free mice, showed that the textile-EDB maintained all animals germ-free, even after 20 droplet sprays at 60cm, while mice located at 180cm became colonized by bacteria-carrying droplets with a single spray (0/8 vs 4/4, Fisher’s exact, p=0.002). Collectively, barriers protected all mice (even with low textile density; heavy vs light fabric, paired t-test, p=0.002) against high droplet doses (2 or 20 sprays) if the EDB fully covered the cage (0/20 vs 10/10, Fisher’s exact, p<0.0001). An overview of the experiment, methods and results is presented in **Figure 1** and **Supplementary Figures 1–2**).

**Figure 1.**
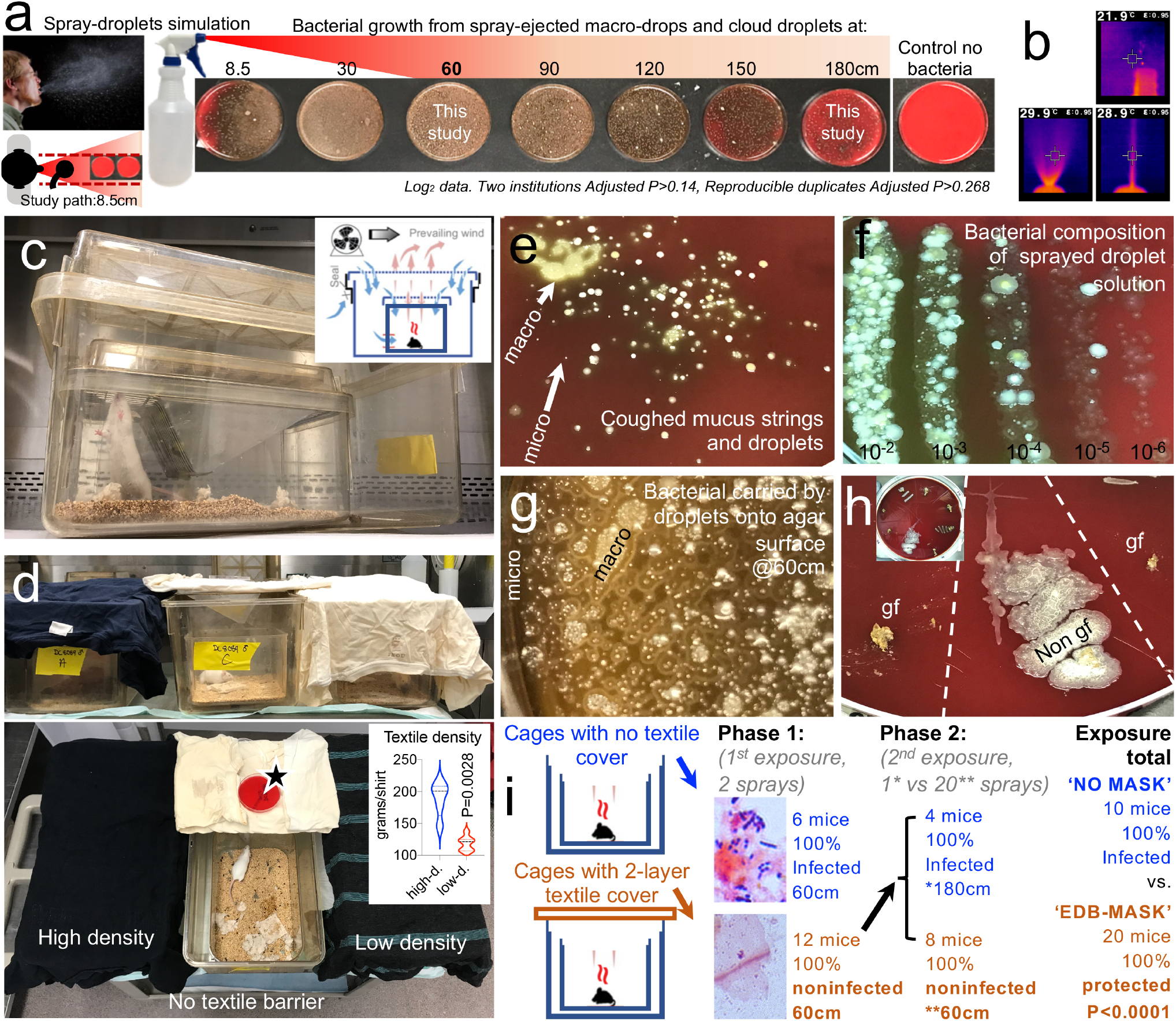
A two-layer textile barrier fully protects germ-free mice from colonization by bacteria contained in sprayed micro-droplets. **a**) Spray-droplet simulation model using bacterial aqueous solution recently validated for the assessment of textiles against COVID-19 in our laboratory. Unmodified from by Rodriguez-Palacios *et al.*^*11*^; open access license. Inset, mechanism of passive filtration. **b**) Thermographic features of cloud-droplet ejection. **c**) Nested isolation caging 2-layer system used to raise germ-free mice. **d**) In this experiment, the two cage lids were replaced by a two-layer textile barrier to prevent EnvDC within the cage. Compared with cages without lid (no mask, surrogate). Sprayed from 60 and 180cm distance. **e**) Visualization of bacteria present in cough microdroplets, healthy adult. TSA plates, aerobic incubation, 48h. **f**) Visualization of rich community composition for the bacteria present in the microdroplet solution used to spray germ-free mice. Parallel lanes plating method^13^. **g**) Visualization of bacteria-contained on macro/micro-droplets on a quarter of a Petri dish. TSA, 21mm horizontal field. **h**) Example of fecal culture-negative from mice protected with textile-EDBs, which remained germ-free (gf), and culture-positive from mice not protected with textile (Non gf), Inset, 20cm plate, 8 samples. **i**) Overview of experiment, results, and fecal gram-stain confirmation (details in **Supplementary Materials**)

## DISCUSSION

As illustrated though the media, the ongoing increase in coronavirus cases has “sparked a ‘war of masks’ in desperate global scramble for protection”15. Despite the seriousness of the mask supply shortage, global institutions have not promoted the mandatory use of homemade masks to prevent COVID expansion and to simultaneously alleviate pressure on medical-grade supplies.

As the main measure to control COVID transmission, virtually everyone in all continents has been requested to ‘stay-at-home’ by lawful orders, and enforcement. Despite such unprecedented, effective global initiative, combined with social distancing (1.8m) as preventive behavior, it is expected that indefinite quarantine may not be sustainable, especially within highly populated and poor regions (currently in their pre-pandemic curve phase). Our results with a single spray towards mice located at 1.8m showed that 100% of mice can get contaminated if not protected with a textile-barrier/mask.

To date, masks have been studied primarily in health care settings and under conditions that are not as publicized or feared as the consequences of the COVID pandemic. Transmitted primarily by oral-respiratory droplets, the COVID-19 pandemic would benefit if the scientists, policymakers, medical advisors, and community have further scientific data to demonstrate that masks are effective to prevent droplet dispersion, while fully protecting individuals from exposure to microbial agents present in the droplets, if masks are properly worn.

This brief report illustrates that germ-free animals when protected by two layers of textile (100% combed cotton, simulating medical mask protection) are fully protected from becoming contaminated with the germs present with a simulated cloud model of bacteria-carrying droplets. In this context, although several studies have shown that masks are effective preventing respiratory infections in humans, masks often fail because often 50% of times people in such settings do not wear them properly^16,17^. This study supports the effective prevention potential of homemade masks rapidly fabricated using widely available cotton fabrics^18^. The U.S. Centers for Disease Control now provides guidance for sewn and non-sewn versions. In addition, the U.S. Surgeon General released a 45-second video with his own tutorial^19^. A mandatory recommendation to wear EDB-textile masks at a global scale will effectively help protect individuals from COVID droplets.

## Availability of data and materials

The raw data supporting the conclusions of this article will be made available by the authors, without undue reservation.

## Competing interests

The authors declare that they have no competing interests.

## Funding

This study was conducted with discretionary funds allocated to ARP.

## Author contributions

ARP envisioned, planned and executed the experiments, analyzed the data, prepared figures and wrote the manuscript. MC, assisted with experiments. FC commented, revised and edited the manuscript. All authors approved the final manuscript.

## Acknowledgements

Special thanks to the Animal Resource Center at Case Western Reserve University and Gina Ponzani for assistance with animal husbandry; Dr. Abigail Basson for proofreading and editorial/scientific suggestions; and Drs. Minh Lam, W. John Durfee, and Kristie Brock for assistance with animal utilization protocol and advice relevant to animal housing and study authorization by IACUC.

---

**Supplementary Figure 1.**
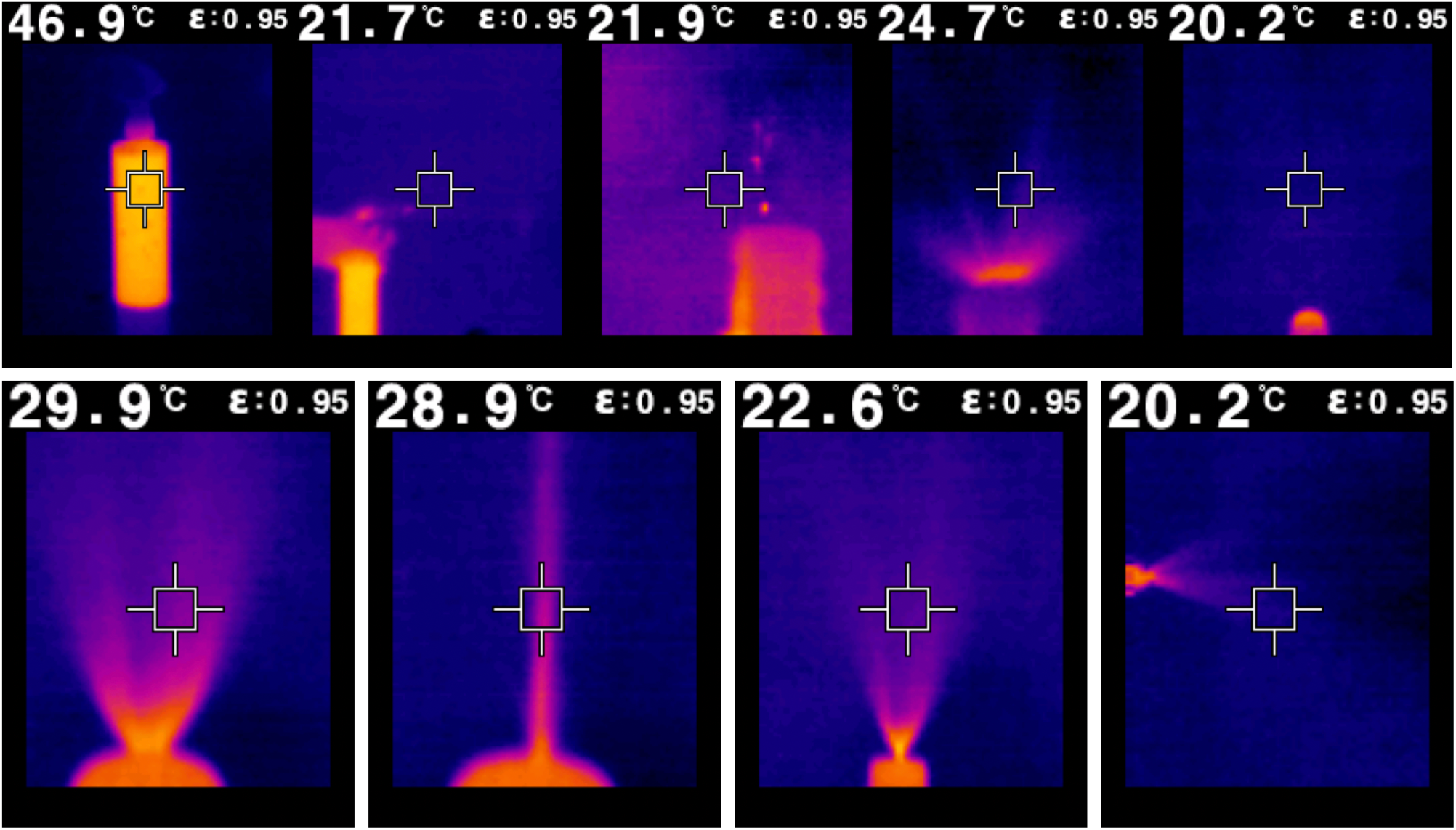
Thermographic characterization of ejection features of our spray macro- and micro-droplet model. Notice the warm liquid solution used (at 46.9°C, rapidly cools down upon ejection as spray. Also note that the complexity of our simulation model resembles the features of the sneeze fluid dynamics as described by Bourouiba *el at.*^*1*^, with wide dispersion of high-velocity microdroplets, splashing of large heavy macro-droplets, and long range projectile-like jet, covering a large conical surface for cloud surface contamination.

**Supplementary Figure 2.**
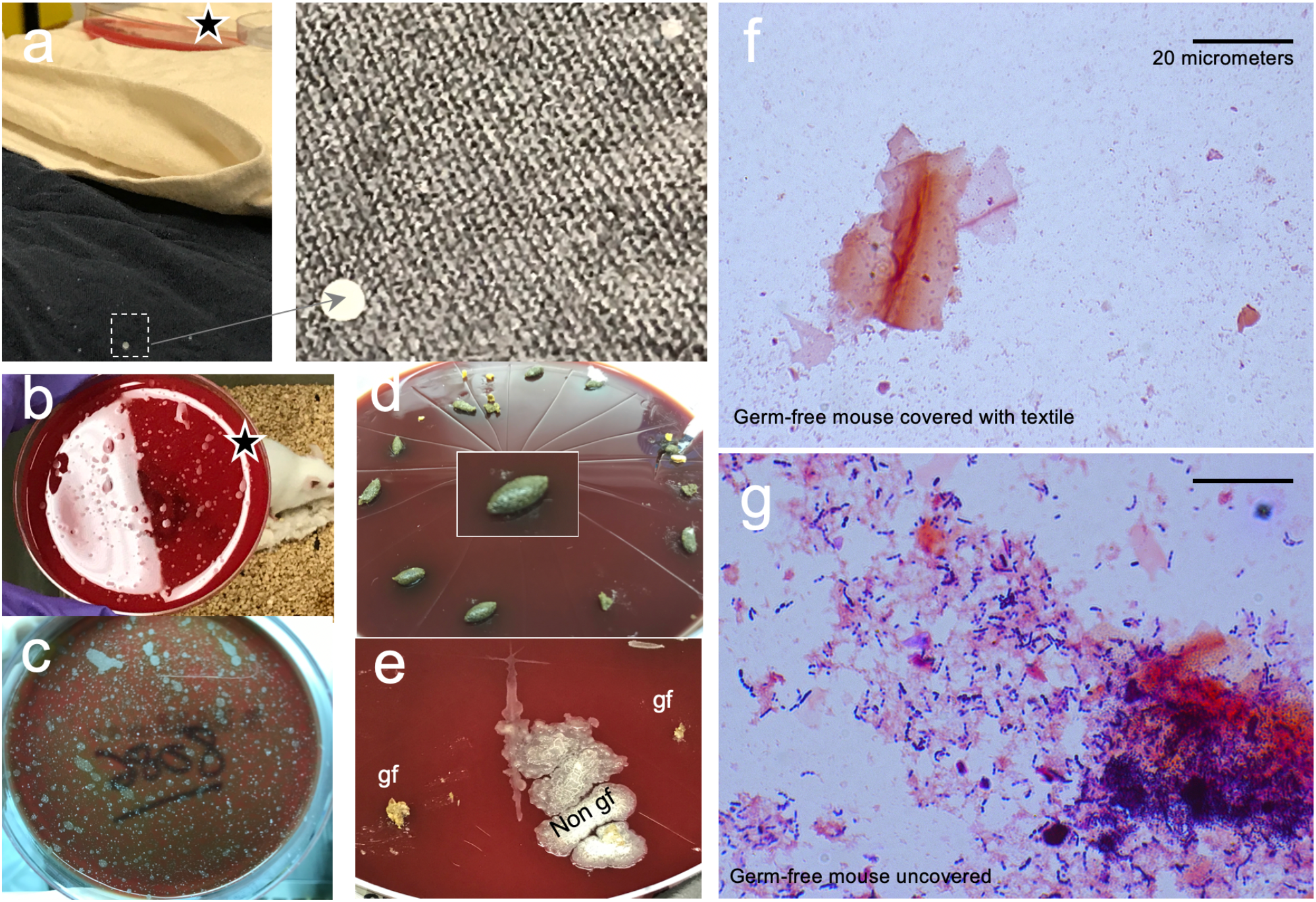
Textile Droplet Barrier fully protect germ-free mice from microbial colonization by bacteria present in sprayed liquid micro-droplets. **a**) Textiles were able to retail large drops and microdroplets. **b**) Agar plate shows droplet density to which mice were exposed immediately after spray (stars). **c**) Aerobic incubation of agar illustrates environmental contamination of mouse cage with numerous microdroplets not visualized immediately after spray. **d**) Fecal samples from all mice show no bacterial growth after 36h of incubation on agar before experiment confirming Germ-free status of mice. **e**) Fecal samples from representative mice protected with textile showing no bacterial growth after 36h of incubation on agar after spray confirming Germ-free status protection by the textile, and no-barrier mice showing fecal bacterial growth. **f-g**) Representative gram stain of fecal samples in this study shown as insets in **Figure 1i**.

## REFERENCES

1 Bedford, T. et al. Cryptic transmission of SARS-CoV-2 in Washington State. medRxiv 2020.04.02.20051417; doi: https://doi.org/10.1101/2020.04.02.20051417 (2020).

2 Mansor, S. Data Suggests Many New York City Neighborhoods Hardest Hit by COVID-19 Are Also Low-Income Areas. Time. Available at: https://time.com/5815820/data-new-york-low-income-neighborhoods-coronavirus/. April 5, 2020. (2020).

3 Dwyer, C. & Aubrey, A. CDC Now Recommends Americans Consider Wearing Cloth Face Coverings In Public. NPR. Coronavirus crisis. Updated. April 3, 2020. Available at: https://www.npr.org/sections/coronavirus-live-updates/2020/04/03/826219824/president-trump-says-cdc-now-recommends-americans-wear-cloth-masks-in-public (2020).

4 Howard, J. Masks may actually increase your coronavirus risk if worn improperly, surgeon general warns. CNN. Updated. March 2, 2020. Available at: https://www.cnn.com/2020/03/02/health/surgeon-general-coronavirus-masks-risk-trnd/index.html. (2020).

5 Howard, J. WHO stands by recommendation to not wear masks if you are not sick or not caring for someone who is sick. CNN. Updated. March 31, 2020. Available at: https://www.cnn.com/2020/03/30/world/coronavirus-who-masks-recommendation-trnd/index.html. (2020).

6 Purohit, K. India COVID-19 lockdown means no food or work for rural poorMillions in underdeveloped regions face penury and deprivation as economic activity grinds to a halt due to lockdown. News/poverty and development. Available at: https://www.aljazeera.com/news/2020/04/india-covid-19-lockdown-means-food-work-rural-poor-200402052048439.html. 3 Apr 2020. (2020).

7 Asadi, S. et al. Aerosol emission and superemission during human speech increase with voice loudness. Sci Rep 9, 2348, doi:10.1038/s41598-019-38808-z (2019).

8 Borowicz, S. A., Plinke, E. & Cahill, T. Ohio Department Of Health Orders Cancellation of All Non-Essential Surgeries to Preserve PPE. Wednesday, March 18, 2020. Available at https://www.natlawreview.com/article/ohio-department-health-orders-cancellation-all-non-essential-surgeries-to-preserve. Accessed March 19, 2020. (2020).

9 WHO. World Health Organization. Coronavirus disease (COVID-19) advice for the public: When and how to use masks. Available at: https://www.who.int/emergencies/diseases/novel-coronavirus-2019/advice-for-public/when-and-how-to-use-masks. Accessed March 18, 2020. (2020).

10 BBC. Coronavirus: Trump to defy ‘voluntary’ advice for Americans to wear masks. BBC. Updated. April 4, 2020. Available at: https://www.bbc.com/news/world-us-canada-52161529. (2020).

11 Rodriguez-Palacios, A., Cominelli, C., Basson, A., Pizarro, T. & Ilic, S. Textile Masks and Surface Covers – A ‘Universal Droplet Reduction Model’ Against COVID-19 Respiratory Pandemic. Submitted to medrxiv March 29, 2020. DOI pending. (2020).

12 Liu, Y. et al. Aerodynamic Characteristics and RNA Concentration of SARS-CoV-2 Aerosol in Wuhan Hospitals during COVID-19 Outbreak/. bioRxiv 2020.03.08.982637; doi: https://doi.org/10.1101/2020.03.08.982637 (2020).

13 Rodriguez-Palacios, A. et al. The artificial sweetener Splenda promotes gut Proteobacteria, dysbiosis and myeloperoxidase reactivity in Crohn’s disease-like ileitis. Inflammatory Bowel Diseases, Accepted, In press. (2018).

14 Rodriguez-Palacios, A. et al. ‘Cyclical Bias’ in Microbiome Research Revealed by A Portable Germ-Free Housing System Using Nested Isolation. Sci Rep 8, 3801, doi:10.1038/s41598-018-20742-1 (2018).

15 Lister, T., Shukla, S. & Bobille, F. CNN - Coronavirus sparks a ‘war for masks’ in desperate global scramble for protection. Updated 1:43 PM ET, Sat April 4, 2020. Available at https://www.cnn.com/2020/04/04/europe/coronavirus-masks-war-intl/index.html. (2020).

16 MacIntyre, C. R. et al. Face mask use and control of respiratory virus transmission in households. Emerg Infect Dis 15, 233–241, doi:10.3201/eid1502.081167 (2009).

17 MacIntyre, C. R. & Chughtai, A. A. Facemasks for the prevention of infection in healthcare and community settings. BMJ 350>, h694, doi:10.1136/bmj.h694 (2015).

18 CDC. Use of Cloth Face Coverings to Help Slow the Spread of COVID-19. Coronavirus Disease 2019 (COVID-19). April 4, 2020. Uso de cubiertas de tela para la cara para ayudar a desacelerar la propagación del COVID-19. Available at https://www.cdc.gov/coronavirus/2019-ncov/prevent-getting-sick/diy-cloth-face-coverings.html.. (2020).

19 Adams, J. Surgeon General Shows How to Make Your Own Face Covering. Surgeon General Dr. Jerome Adams. Tutorial April 4, 2020. Available at https://www.youtube.com/watch?v=PI1GxNjAjlw. (2020).

